# EvoPRAISE: computationally guided directed evolution for rational AAV capsid engineering

**DOI:** 10.1101/2025.11.02.686002

**Authors:** Hiroaki Ono, Shoko Fujino, Genshiro A. Sunagawa

## Abstract

Advances in neuroscience and gene therapy would be greatly accelerated by adeno-associated virus (AAV) vectors capable of crossing the blood–brain barrier (BBB). Unfortunately, capsids evolved in mice often fail to function in other species, limiting applications in non-model animals such as hibernators. To overcome this limitation, we developed Evolutionary Optimization combined with APPRAISE (EvoPRAISE), a framework that integrates structure-based peptide–protein affinity prediction with directed evolution to design peptide binders for membrane proteins. Using EvoPRAISE, we designed peptide binders that interact with Ly6E, a BBB-associated membrane protein expressed in the Syrian hamster brain. By inserting these peptides into surface-exposed regions of the AAV9 capsid, we developed AAV capsid variants capable of crossing the BBB following systemic administration in Syrian hamsters. Furthermore, this framework enabled the design of AAV capsids displaying peptide binders targeting the human BBB. This rational approach reduces the need for large-scale animal *in vivo* selection procedures and facilitates the efficient development of receptor-targeted AAV capsids for both neuroscience studies in non-model species and translational gene delivery to the human brain.

## INTRODUCTION

Recombinant adeno-associated viruses (rAAVs) are widely used as gene-delivery vectors for basic research and gene therapy. Indeed, rAAVs can transduce both dividing and non-dividing cells, persist predominantly as episomal concatemers with long-term expression, and elicit comparatively low innate and adaptive immune responses^1–4^. rAAV genomes are compact (∼4.7 kb) single-stranded DNA molecules flanked by inverted terminal repeats, which constrains payload size but also enables modular vector design with diverse promoters and cargoes^5,6^. Among naturally occurring serotypes, AAV9 has emerged as a versatile scaffold for engineering tissue-targeted vectors owing to its broad biodistribution after systemic administration and its proven manufacturability^7–12^. Building on this scaffold, variable capsid-engineering strategies have produced variants with enhanced and retargeted tropism^11^. For example, AAV-PHP.B and AAV-PHP.eB were generated to realize efficient, widespread central nervous system (CNS) transduction after intravenous delivery in mice^10,11^. Notably, AAV.Cap-B10 was engineered to maintain strong CNS tropism in mice while minimizing off-target liver transduction^12^.

Much of modern capsid engineering relies on *in vivo* selection of highly diverse libraries (typically 10^5^–10^7^ variants) generated by peptide display (e.g., 7-mer NNK)^10,11^, error-prone mutagenesis^13,14^, and/or DNA shuffling^15–17^. After systemic administration, enriched variants are recovered from target tissues by next-generation sequencing of barcodes or capsid ORFs. Iterative rounds of this procedure yield candidates with improved performance. This directed-evolution paradigm is powerful precisely because it does not require prior knowledge of the target receptor or mechanism of entry. However, AAV variants isolated in one host frequently fail to generalize, even across mouse strains, and seldom translate to non-human primates. For example, PHP.eB crosses the blood–brain barrier (BBB) in C57BL/6 but loses activity in BALB/c, reflecting strain-specific expression of entry factors and vascular contexts^18,19^. More broadly, rodent-selected CNS-tropic capsids have not reproduced robust BBB transduction in primates^20,21^.

These translation gaps become a major problem when a phenotype of interest spans taxa. Hibernation exemplifies this challenge. Hibernating mammals can survive prolonged periods of scarcity by markedly suppressing basal metabolic rate and lowering core body temperature to ∼10 °C, often for days or weeks^22–24^. In laboratory mice, exposure to low ambient temperature and food restriction can elicit daily torpor. This condition is superficially similar to hibernation, albeit shorter and comprising a single-cycle state with distinct thermoregulatory control mechanisms^25,26^. Recent work has identified hypothalamic neuronal populations that can drive hibernation-like hypothermia and hypometabolism in mice, implicating the CNS in the control of these states^27,28^. Yet the extent to which these circuits and their molecular entry points are conserved in natural hibernators remains unclear. Hibernators that can be maintained in laboratory settings (e.g., Syrian hamsters) currently lack fully optimized genetic-modification toolkits. As such, AAV-mediated gene delivery is an attractive approach for CNS studies of hibernation.

In recent years, rational design has become far more accessible thanks to advances in structural biology and the accumulation of domain knowledge^29,30^. Rational design offers a practical route to engineering capsids that translate across species, including humans, without the need for large *in vivo* selection experiments. Using this framework, tissue tropism is programmed by grafting receptor-binding modules (e.g., short peptides or nanobodies) into permissive, surface-exposed loops of the AAV capsid so that the vector engages proteins displayed by the target tissue. Rapid progress in protein-structure informatics, exemplified by AlphaFold2/Multimer, has made *de novo* peptide-binder design increasingly realistic^31–34^. Incorporating these binders into capsid loops enables the computational design of receptor-targeted AAVs without extensive experimental screening. Notably, Automated Pairwise Peptide–Receptor Analysis for Screening Engineered proteins (APPRAISE) evaluates candidate peptide–receptor pairs by building complex models (via AlphaFold-Multimer^35^ or ESMFold^36^) and scoring them with fast, interpretable metrics that incorporate biophysical and geometric constraints^37,38^. Crucially, by considering interface complementarity and steric accessibility, APPRAISE goes beyond simple affinity estimates between peptide and receptor to approximate whether a capsid-displayed peptide could physically access its receptor. This methodology has been validated across multiple protein classes, including miniproteins targeting the severe acute respiratory syndrome coronavirus 2 (SARS-CoV-2) spike, nanobodies targeting a G-protein-coupled receptor, and peptides that bind to the transferrin receptor or programmed death-ligand 1 (PD-L1). Its accuracy, interpretability, and generalizability suggest that APPRAISE can accelerate structure-guided capsid engineering for biomedical applications.

Here, we introduce EvoPRAISE, an *in silico* directed-evolution framework that couples APPRAISE-based ranking with algorithmic diversification. One limitation of APPRAISE is that the output gives relative rankings within a user-specified peptide pool. The absence of a known positive-control binder can make outcomes dependent on the initial pool composition. EvoPRAISE does not rely on a fixed pool but instead iteratively mutates top- ranked sequences to explore new neighborhoods of sequence space. In this way, EvoPRAISE identified variants to improve the physics-informed binding scores defined by APPRAISE. Indeed, the directed-evolution framework minimized pool bias and helped escape local optima. Applied to CNS delivery, EvoPRAISE yielded AAV capsids that cross the BBB and transduce the brain in Syrian hamsters, a laboratory-manageable hibernator, and also identified AAV capsids that bind with proteins expressed at the human BBB. Together, this framework enables rational, cross-species-compatible capsid design while reducing dependence on large-scale *in vivo* experimental screening programs.

## RESULTS

### EvoPRAISE: *in silico* directed evolution method for engineering the AAV capsid

The EvoPRAISE workflow (Figure 1A) comprises three main steps. In the first step, randomly generated peptides are computationally evaluated for their binding score (*B*) to the target protein using APPRAISE^37^. The APPRAISE binding score is a non-negative metric derived from atom–atom contact counts and geometric constraints (i.e., binding angle and pocket depth), which is critical for peptide binding. Based on this binding score, peptides are ranked, and the top-ranking candidates selected. In the second step, the selected top- ranking peptides undergo saturated mutagenesis. Saturation mutagenesis is performed by substituting each residue with one of the remaining 19 commonly occurring natural amino acids, yielding a library of 19*L* single-mutant variants (where *L* is the peptide length in amino acids). In the third step, the resulting mutant library is reevaluated by APPRAISE, and the top ranked binders selected. By iteratively repeating steps two and three, peptides are progressively optimized toward higher binding affinity with the target protein, following the principles of directed evolution.

**Figure 1.**
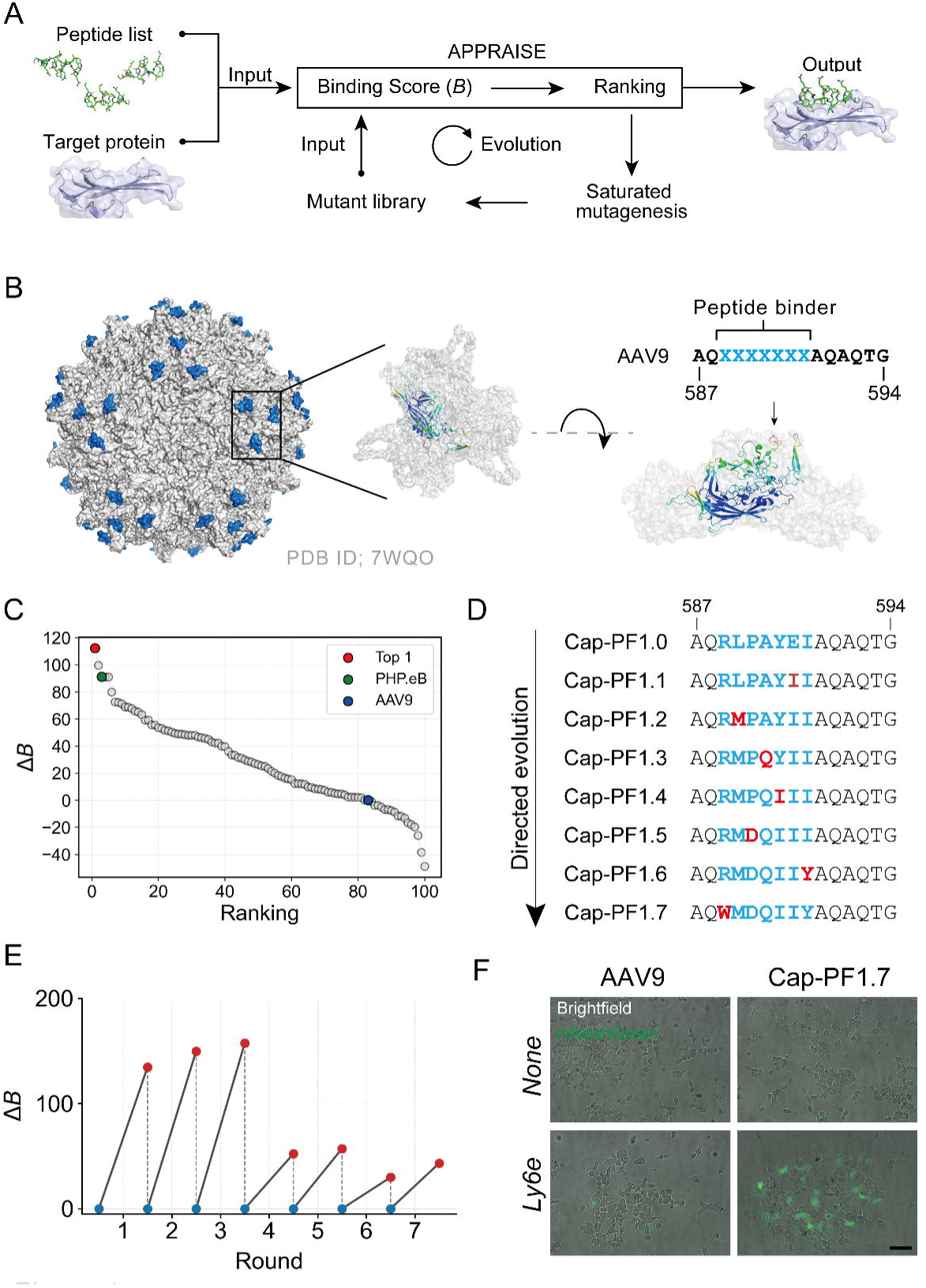
Workflow of EvoPRAISE and its application in generating a peptide binder for Ly6e (A) Workflow of EvoPRAISE. Firstly, the extracellular domain of the target membrane protein and a pool of randomly generated peptides were prepared. Secondly, these inputs were processed by APPRAISE, which calculates an energetic binding score (B) for each peptide based on atom counting. Peptides were ranked according to binding score, and the top candidate was selected (Round 0). For the top-ranked peptide, a saturation mutagenesis library was generated by substituting each residue with one of the 19 other common natural amino acids. This library was then evaluated by APPRAISE to determine a new top-ranking peptide (Round 1). In subsequent rounds, residues that had already evolved were fixed, while saturation mutagenesis was applied to the remaining residues. This iterative process was repeated until all residues had been evolved. (B) Structural model of AAV-PHP.eB, which highlights the peptide insertion site in blue. The left panel shows the AAV capsid composed of 60 structurally identical subunits (PDB ID: 7WQO). The middle panels show top views around the threefold symmetry axis, with the three subunits forming the trimer displayed. A single VP3 subunit is highlighted in green, and the inserted peptide sequence is shown in blue. Peptide sequence used as the EvoPRAISE input. Seven-residue peptide binders were inserted between residues 588 and 589 (VP1 numbering) in a surface-exposed variable region of AAV9. (C) Binding scores of 100 randomly generated peptides were compared with that of the AAV9 peptide (AQAQAQTG) and plotted as ΔB in ranking order. In Round 0, the RLPAYEI peptide (red) ranked first. The peptide pool also included the PHP.eB peptide (green) and the AAV9 peptide (blue). (D) Changes in binding scores of the top-ranked peptides across rounds. Red plots indicate the top peptide of each round. In the subsequent round, the same peptide was used as the reference for comparison against its variants (blue). (E) *In vitro* infectivity assay. AAV-Cap-PF1.7 showed Ly6e-dependent enhancement of transduction in HEK293T LY6E-KO cells overexpressing Ly6e, whereas the negative-control AAV9 did not. AAV capsids carrying a fluorescent protein expression cassette were applied at 5 × 10^9^ viral genomes (v.g.) per well to HEK293T cells transfected or not with Ly6e in a 96- well plate format. Images were taken 24 h after transduction (n = 3 per condition). Scale bars, 200 µm.

We first applied EvoPRAISE to engineer rAAV capsids capable of targeting the brain. Previous large-scale *in vivo* screening studies have yielded variants such as AAV- PHP.eB^39^ and AAV.Cap-B10^12^, which can cross the BBB and efficiently transduce the brain. These capsids are AAV9-derived variants displaying short (7 amino acid) surface peptides. Genetic and biophysical studies have shown that binding to LY6A, a GPI-anchored membrane protein, is essential for BBB penetration and CNS entry^40^. As a demonstration of EvoPRAISE, we aimed to design peptide binders for the Ly6 family expressed in the Syrian hamster brain, with the goal of generating an AAV capsid capable of transducing the CNS of the Syrian hamster. We designed 7-mer peptide binders to be inserted into residues 588– 589 of the VP1 capsid protein, a surface-exposed region that can directly contact receptors in the assembled capsid^41,42^ (Figure 1B).

To determine the molecular target, we examined Ly6 family genes expressed in the Syrian hamster brain. Genome analysis (GCF_017639785.1 assembly) identified at least 18 Ly6 family genes (Figure S1A). To select the most abundantly expressed transcript, qPCR was performed using bulk brain samples, which revealed that *Ly6e* was the most highly expressed gene (Figure S1B). We therefore set Ly6e as the target protein for EvoPRAISE.

As a starting point for the directed evolution, 100 random 7-mer peptides were generated along with two capsid controls (PHP.eB and wild-type AAV9). The extracellular domain of Ly6e (residues 27–118) was used as the target receptor input. APPRAISE ranked the peptides according to their binding score to Ly6e (residues 27–118). This analysis revealed the RLPAYEI peptide (Cap-PF1.0) to display the top binding score (Figure 1C). Next, directed evolution of Cap-PF1.0 was initiated. A mutant library in which each amino acid of RLPAYEI was substituted with all other 19 natural amino acids was generated. Because exhaustive evaluation of the full single-mutant library in one run would require several days of computation even on an NVIDIA V100 GPU (Google Colab), we applied APPRAISE to identify the top-ranked mutant at each amino-acid position. The best candidates from all positions were then pooled, and the top overall binder (RLPAYII: Cap-PF1.1) selected. Thus, Cap-PF1.1 was used as the seed for the next round of directed evolution. In each round, once a position yielded a beneficial substitution, it was fixed and excluded from further randomization. This lock-in strategy ensured that all seven positions were systematically optimized over seven rounds. At each round, the APPRAISE binding score for the lead peptide exceeded that of the previous round, yielding stepwise gains in predicted Ly6e binding (Figure 1D-E). Across the Cap-PF1.0→PF1.7 trajectory, the seven-residue insert evolved from RLPAYEI to WMDQIIY, showing alternating shifts in hydropathy (grand average of hydropathy (GRAVY))^43^ and net charge (Figure S2). The initial two rounds increased hydrophobicity by replacing a negatively charged residue (E→I) and a hydrophobic residue with a bulkier hydrophobic amino acid (L→M), setting the net charge to +1. A polar substitution (A→Q) then reduced the GRAVY value without affecting the charge, followed by Y→I, which restored hydrophobicity. Introduction of a negative charge (P→D) dropped the net charge from +1 to 0 and decreased the GRAVY value. The substitution I→Y further lowered hydrophobicity while leaving the charge unaltered. Finally, R→W shifted the net charge to −1 while partially restoring hydrophobicity. Thus, overall, the net charge progressed 0 → +1 → 0 → −1, and hydrophobicity oscillated accordingly, revealing a compensate-and-balance pattern that converged on Cap-PF1.7 (WMDQIIY).

The final evolved variant peptide WMDQIIY was named Cap-PF1.7. To validate the interaction between Ly6e and Cap-PF1.7, *in vitro* potency assays^40^ were performed. HEK293T cells were transiently transfected with plasmids encoding the Syrian hamster Ly6e and the effects on binding and transduction by Cap-PF1.7 capsids evaluated. Because Ly6e has a human ortholog (LY6E), we generated LY6E-knockout HEK293T cells to prevent cross-reactivity. Under these conditions, Cap-PF1.7 showed markedly increased binding and transduction in a Ly6e-dependent manner (Figure 1E), supporting a role for Ly6e in Cap- PF1.7–mediated entry.

### The evolved AAV capsid variant crosses the BBB of the Syrian hamster

To validate the *in silico* predictions, we assessed the tissue tropism of Cap-PF1.7 after systemic administration in Syrian hamsters. For comparison with capsids developed for CNS tropism in mice, AAV9, AAV-PHP.eB, AAV-Cap.B10, and Cap-PF1.7 were used to package a reporter in which mNeonGreen was expressed under the ubiquitous CAG promoter^39^ (Figure 2A). All capsids produced virus with comparable efficiencies (Figure 2B), indicating that Cap-PF1.7 did not impair AAV capsid fitness. In each case, 1 × 10^12^ vector genomes (v.g.) of AAV variants were intravenously administered to 3-week-old hamsters. The hamsters were weaned and large enough to permit retro-orbital injection into the retro-orbital- sinus (behind the globe of the eye). Transduction efficiency was assessed 4 weeks post-injection by monitoring mNeonGreen expression. As a result, AAV9, AAV-PHP.eB, and AAV- Cap.B10 failed to transduce the Syrian hamster brain, whereas Cap-PF1.7 efficiently transduced the adult CNS, with broad mNeonGreen expression in the cortex, hippocampus, and thalamus (Figure 2C). Moreover, strong tropism for fiber tracts, such as the striatum and the dorsal part of the spinal cord (spinal trigeminal nucleus), was also observed (Figure 2D). To determine the cell types transduced by AAV-Cap-PF1.7, we performed Nissl staining in brain regions that exhibited strong transduction. Quantitative analysis revealed that mNeonGreen expression colocalized with Nissl-positive neurons in 91.6% of cells in the hippocampus, 70.7% in the cortex, and 93.3% in the thalamus (Figure S3A). These results indicate that neurons are the principal cellular targets of AAV-Cap-PF1.7 in the hamster brain. To assess potential off-target infection, we examined mNeonGreen expression in peripheral tissues and found that AAV-Cap-PF1.7 showed limited off-target infection in the liver (Figure S3B).

**Figure 2.**
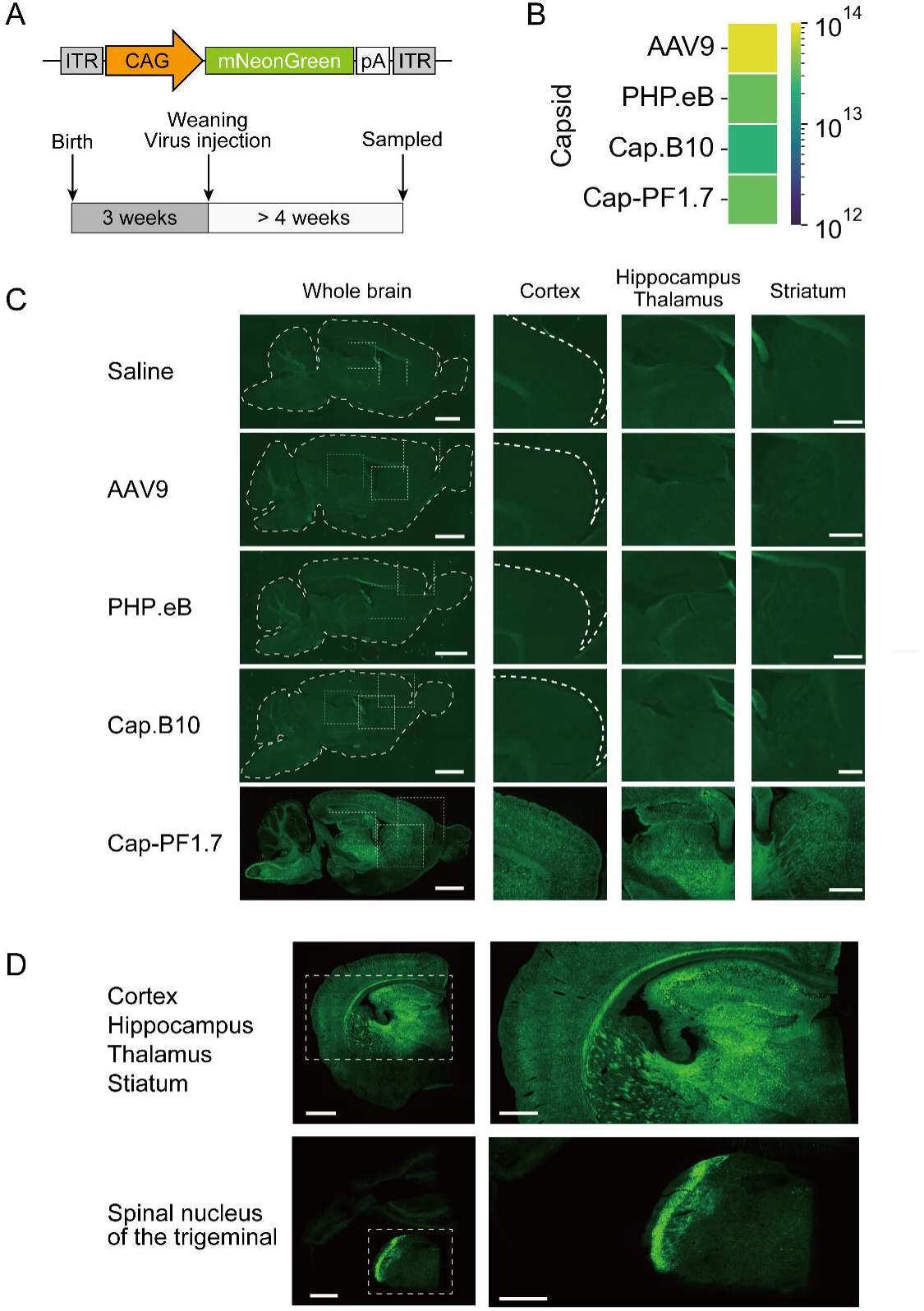
AAV.Cap-PF1.7 crosses the blood-brain barrier (BBB) of the Syrian hamster (A) Experimental scheme to assess capsid tropism for the central nervous system (CNS) *in vivo*. The AAV genome was engineered to express mNeonGreen under the control of a CAG promoter. Vectors were administered systemically to weaning Syrian hamsters via the retro- orbital venous plexus. Animals were sampled ≥4 weeks post-injection, and mNeonGreen expression in the CNS was examined by fluorescence microscopy. (B) Evaluation of the impact of peptide insertion on capsid fitness based on production yield (v.g./mL/20 mm dish). The color bar indicates the absolute mean yield (n = 2–4 per variant). (C) Fluorescence imaging of Syrian hamster brains. AAV9, AAV-PHP.eB, Cap.B10, and Cap-PF1.7 packaging CAG–mNeonGreen were administered intravenously at 1 × 10^12^ v.g. per animal (n = 3 per condition). (D) Representative sections from Cap-PF1.7–injected animals: a coronal brain section at the striatal level and a transverse section at the dorsal spinal cord level. Scale bars, 3 mm (sagittal and coronal image) and 500 µm (brain region).

These results demonstrate that AAV-Cap-PF1.7 is a novel serotype capable of crossing the BBB in the Syrian hamsters and mediating widespread neuronal transduction throughout the CNS.

### Changes in tropism throughout the evolution process

We leveraged EvoPRAISE’s ability to track sequence trajectories to ask how CNS tropism emerges during *in silico*–guided evolution. To this end, pAAV-CAG- mNeonGreen was packaged into capsids displaying each peptide along the Cap-PF1.x trajectory and CNS transduction evaluated under the same experimental conditions as shown in Figure 2A. Across the series (Cap-PF1.0 to Cap-PF1.7), all variants yielded comparable vector production, indicating that the mutations introduced during evolution did not compromise capsid fitness (Figure 3A).

**Figure 3.**
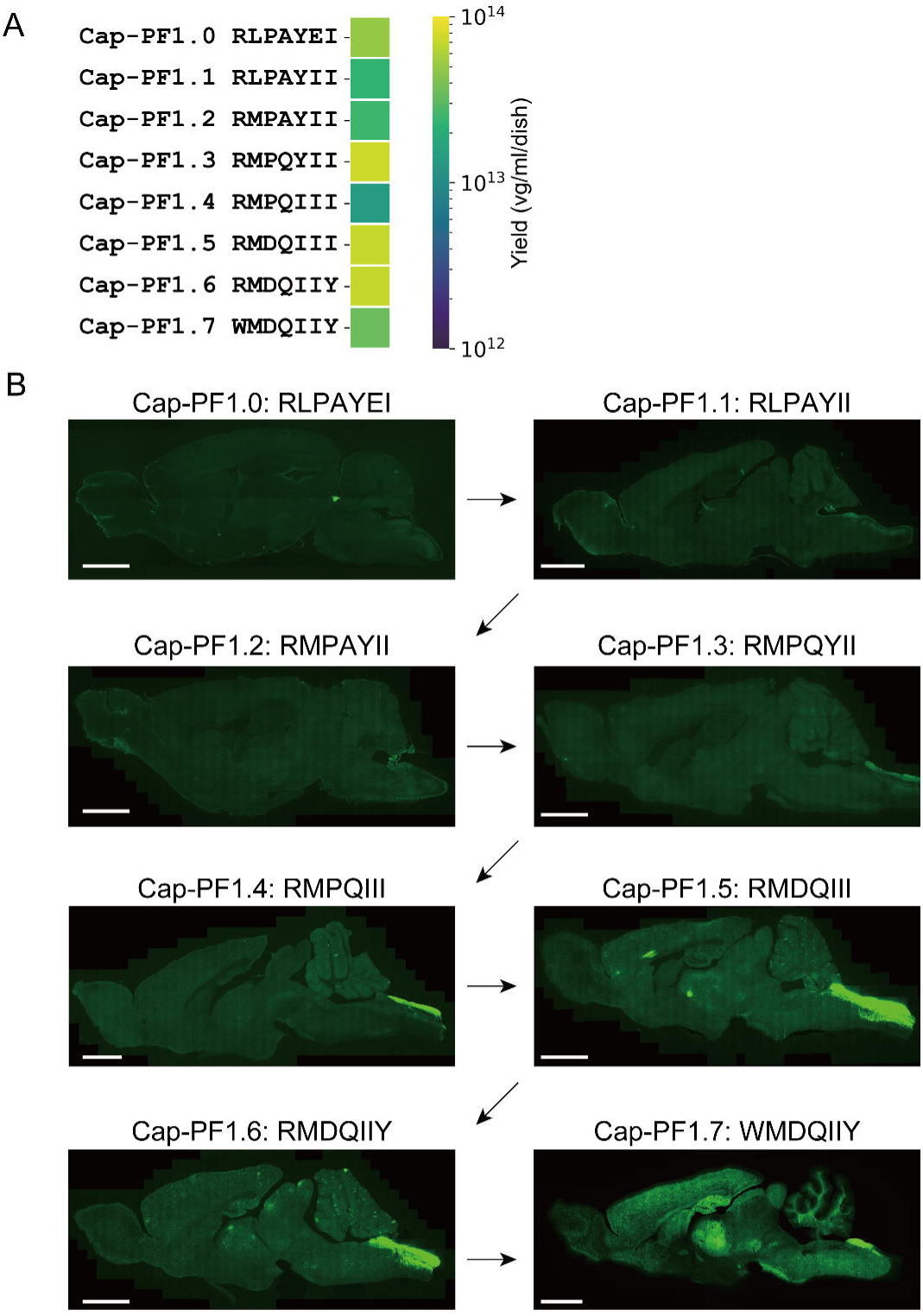
Directed evolution enhances the infectivity of AAV-CapPF1.7 within the central nervous system (CNS) (A) Evaluation of the impact of peptide insertion on capsid fitness based on production yield (v.g./mL/20 mm dish). The color bar indicates the absolute mean yield (n = 2–4 per variant). (B) Representative peptide-binder capsids from each round of directed evolution were used to package mNeonGreen under the control of the ubiquitous CAG promoter. Vector genomes were administered intravenously to Syrian hamsters at a dose of 1 × 10^12^ v.g. per animal (n = 3 per condition). Four weeks after administration, transgene expression was assessed by visualizing mNeonGreen fluorescence throughout the brain, demonstrating that the directed evolution process increased potency. Scale bars, 3 mm.

Cap-PF1.0, the top-ranked APPRAISE binder from an initial pool of 100 random peptides, displayed no detectable mNeonGreen signal in the CNS. Similarly, Cap-PF1.1 and Cap-PF1.2 showed no measurable transduction. A faint but spatially restricted signal in the dorsal spinal cord was evident for Cap-PF1.3, which was more intense for Cap-PF1.4. Cap- PF1.5 showed robust parenchymal transduction in multiple brain regions, including cortex, thalamus and hippocampus, which was further intensified for Cap-PF1.6. The transition from Cap-PF1.6 to Cap-PF1.7 produced a marked, brain-wide increase in transduction, consistent with a late-stage consolidation of CNS tropism across regions (Figure 3B). Thus, APPRAISE serves as a useful tool when the initial pool contains promising peptide binders or positive controls. However, when prior knowledge concerning peptide–receptor interactions is scarce, as in the case of targeting non-model animals, expanding the search space through iterative diversification with EvoPRAISE is advantageous.

### Interaction of the VP3 trimer complex of Cap-PF1.7 with Ly6e

We next interrogated the structural basis of the Cap-PF1.7–Ly6e interaction in a two-step process. Firstly, the candidate binding mode for the Cap-PF1.7–Ly6e complex was generated and evaluated computationally. Secondly, experimental validation was performed using a cell-based potency assay.

For the computational analysis, the Cap-PF1.7–Ly6e binding mode was analyzed with AlphaFold3-Multimer^30^. In this model, the peptide fits into the globular extracellular domain of Ly6e (Figure 4A). Within Cap-PF1.7, Trp1’ and Met2’ were identified as high-confidence positions using the <u>p</u>redicted local <u>d</u>istance <u>d</u>ifference test (pLDDT) metric.

**Figure 4.**
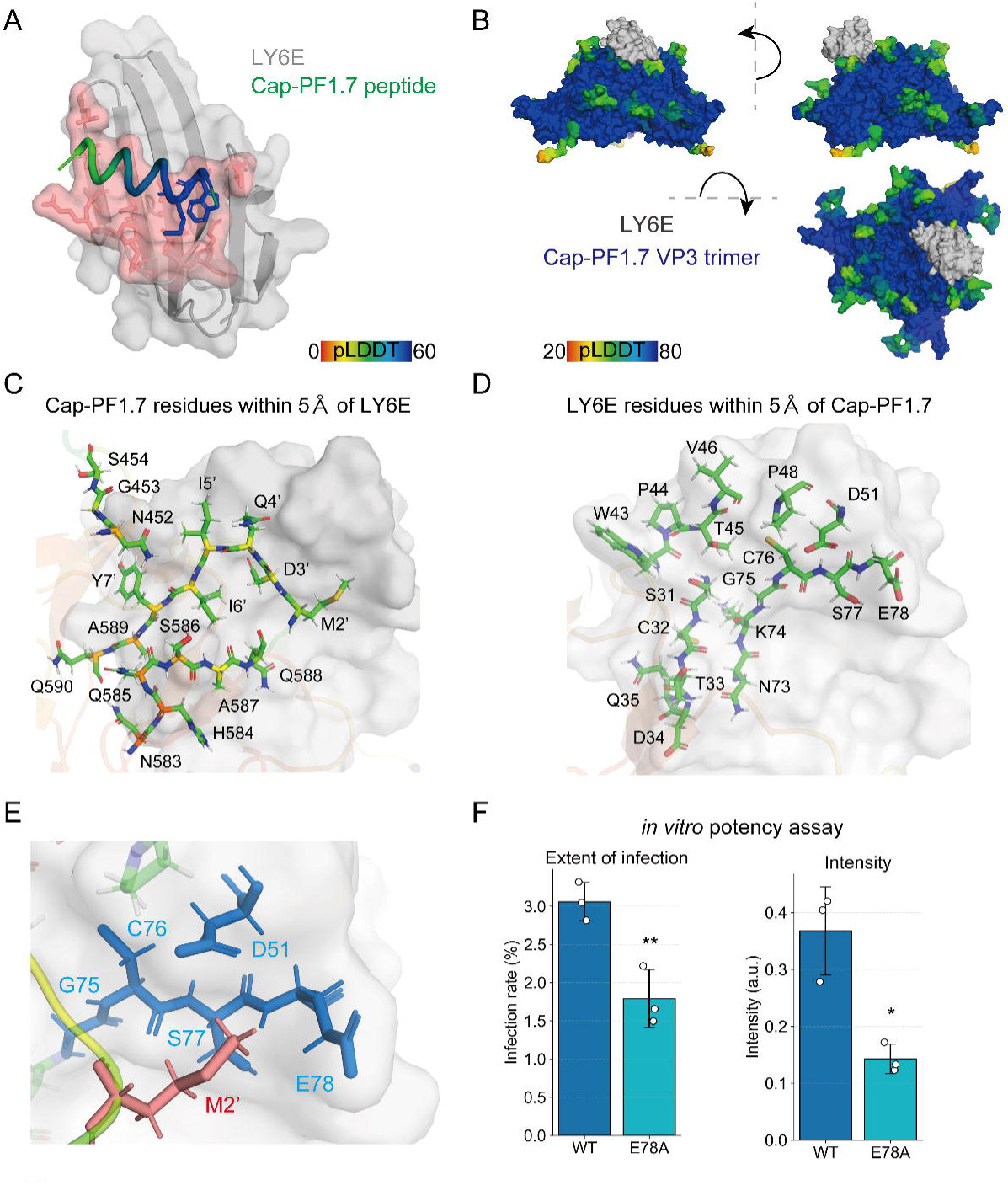
Structural analyses of the binding mode between AAV.Cap-PF1.7 and Ly6e (A) AlphaFold3-Multimer–predicted complex of Ly6e with the Cap-PF1.7 peptide. The peptide is color-coded by per-residue model confidence (i.e., pLDDT score). Ly6e residues containing ≥2 atoms within 5 Å of any peptide atom are highlighted in red and defined as interface residues. High-confidence residues Trp1’and Met2’of Cap-PF1.7 are shown as sticks. (B) Overall binding mode of the Cap-PF1.7–bearing VP3 trimer in complex with Ly6e. The AlphaFold3-Multimer prediction from (A) was integrated with the Cap-PF1.7 VP3 trimer model, and loop structures at the binding interface were refined using RosettaRemodel within the AAV trimeric context. The final model was further optimized using the Rosetta fast-relax protocol. (C) Close-up view of the Cap-PF1.7–Ly6e binding interface. Cap-PF1.7 residues containing ≥2 atoms within 5 Å of Ly6e are highlighted. (D) Close-up view of the Cap-PF1.7– Ly6e binding interface from the Ly6e side. Ly6e residues containing ≥2 atoms within 5 Å of Cap-PF1.7 are highlighted. (E) Interface detail showing Ly6e residues (blue) with ≥2 atoms within 5 Å of Met2’of the Cap-PF1.7 peptide (red). The VP3 loop structure is shown in yellow. (F) Functional validation of the interface by Ly6e mutagenesis. Substitution of the top-contact residue (E78A) reduced potency in a cell-culture infectivity assay. Left, percentage of infected cells (n = 3). Right, total mNeonGreen signal intensity per positive area (n = 3). Error bars indicate mean ± s.d.; p < 0.05, p < 0.01 by Student’s t-test.

To incorporate the receptor interaction into the native capsid context, the Cap- PF1.7 peptide and Ly6e were modeled on the AAV threefold-symmetry spike. Because the AAV VP3 trimer (∼200 kDa) is a large protein complex, AlphaFold3-Multimer did not capture direct contacts between Ly6e and the VP3 trimer bearing Cap-PF1.7. Therefore an integrative structural modeling pipeline was employed similar to that described by Shay et al^38^. Firstly, the AlphaFold-predicted AAV VP3 trimer was structurally aligned with the AlphaFold- predicted receptor–peptide complex through the high-confidence Trp1’ and Met2’ residues of Cap-PF1.7. Secondly, to refine the geometry of the inserted loop, the entire Cap-PF1.7 sequence was optimized using RosettaRemodel^44^, with Ala587 and Ala589 (VP1 indices) serving as anchor residues. Finally, the resulting models were further refined using the Rosetta fast-relax protocol to minimize total energy and improve local structural accuracy (Figure 4B). To identify the interface between the VP3 trimer and Ly6e, we defined contacting residues as those containing at least two atoms within 5 Å of each other. The final model revealed 17 capsid residues at the Ly6e interface, including all amino acids of the Cap-PF1.7 peptide except Trp1’ (Figure 4C). Conversely, 17 Ly6e residues were found in contact with the VP3 trimer (Figure 4D). Based on this structural analysis, Met2’ of Cap-PF1.7 showed a high pLDDT score and multiple contacts with Ly6e residues. These findings suggested Met2’ plays an important role in generating the interacting surface. Among the five Ly6e residues contacting Met2’, Glu78 exhibited the highest number of atomic contacts (Figure 4E).

To test the functional relevance of this interface an E78A substitution was introduced in Ly6e and an *in vitro* potency assay performed. The mutation markedly reduced both the infection rate and mNeonGreen signal intensity compared with wild-type Ly6e, supporting the predicted binding mode and implicating Glu78 as a key determinant of the Cap-PF1.7–Ly6e interaction (Figure 4F).

### An AAV capsid variant that interacts with human Ly6e

Finally, as an application of EvoPRAISE, peptides were designed that target human LY6E. AAV capsids capable of crossing the human BBB would represent a major advance for noninvasive gene therapy, particularly for neurological diseases such as Parkinson’s disease^45^. To assess *LY6E* expression in the human brain the Human Protein Atlas^46–48^ was consulted. *LY6E* mRNA levels across brain regions were compared with BBB marker genes *TFRC* and *SLC2A1*. *LY6E* was found to be expressed at higher levels than *TFRC* in all regions and at levels comparable to or exceeding *SLC2A1* in many regions (Figure 5A). These findings suggested that AAV capsids capable of binding LY6E might have the potential to cross the human BBB. Ten candidate LY6E-binding variants were generated using EvoPRAISE. Cap-PF.h-9 and Cap-PF.h-10 were excluded from further assays due to low AAV yields, consistent with impaired capsid fitness and/or structural destabilization (Figure 5B). *In vitro* potency assays on the remaining eight variants showed that nearly all exhibited stronger transduction signals than AAV9 (Figure 5C). These results suggest that, given a defined target receptor, EvoPRAISE provides a generalizable route to engineer AAV capsids with functional receptor interactions across diverse targets.

**Figure 5.**
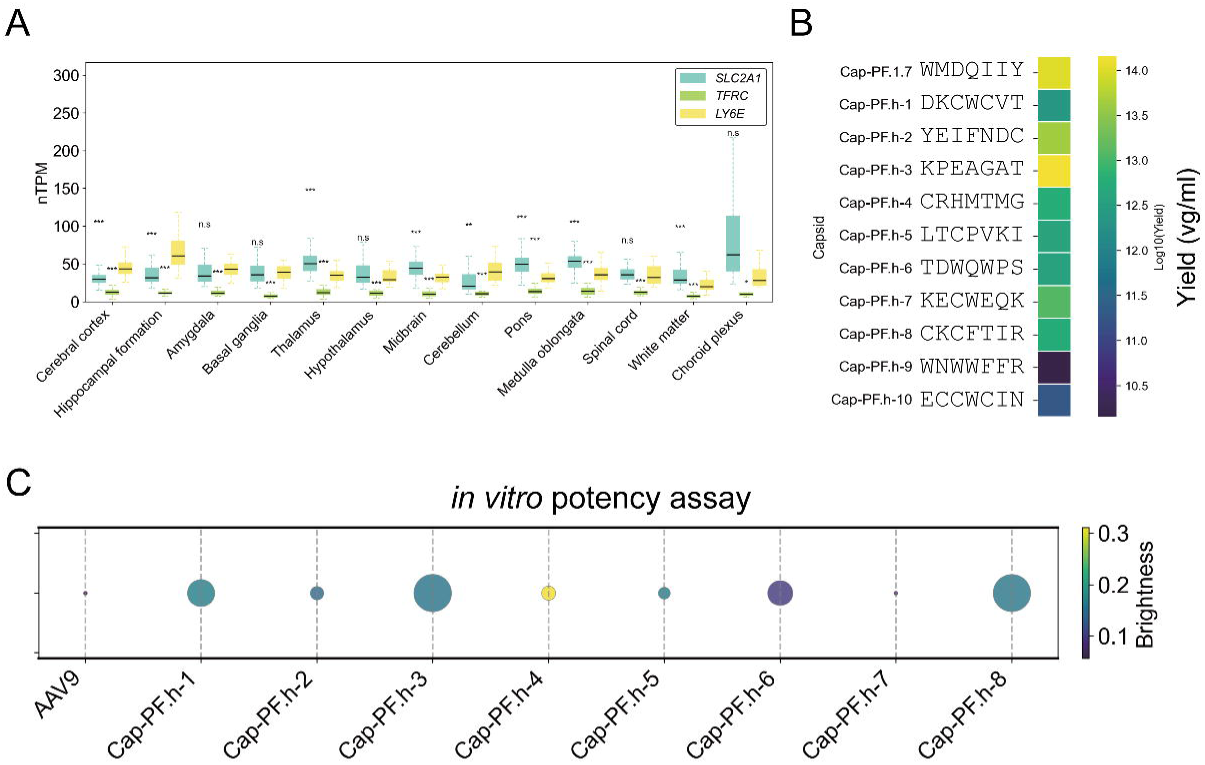
AAV capsid variant that interacts with human LY6E (A) Comparison of *SLC2A1*, *TFRC*, and *LY6E* mRNA expression levels across human brain regions using RNA expression data from the Human Protein Atlas (https://www.proteinatlas.org). *SLC2A1* and *TFRC* are representative marker genes of the blood–brain barrier (BBB). For comparisons between *LY6E* and the other genes within each brain region, the Mann–Whitney U test was applied followed by multiple testing correction using the Benjamini–Hochberg method. Statistical significance is indicated as follows: p < 0.05 (*), p < 0.01 (**), p < 0.001 (***). (B) Impact of peptide insertion on capsid fitness, assessed by production yield (v.g./mL/20 mm dish). The color bar indicates the absolute mean yield (n = 2–4 per variant). (C) Potency of engineered AAVs in HEK293T cells transfected with LY6E, showing extent of infection (max, 1.2; min, 0.01) and total brightness per signal area (max, 0.55; min, 0.055).

## DISCUSSION

Crossing the BBB to deliver genes into the CNS enables mechanistic studies of brain physiology and supports therapeutic development. Many research groups have modified AAV capsids to enhance CNS tropism^10–12,49–52^. In mice, large-scale *in vivo* library screens can identify BBB-penetrant variants, such as AAV-PHP.eB in the C57BL/6 strain. However, translating these findings to non-model species, including humans, remains challenging due to limited cohort sizes and the lack of strain-matched genetic tools. Rational capsid design allows researchers to screen candidates *in silico* instead of directly testing large libraries in animals. Using this strategy limited *in vivo* resources are focused on the most promising cross-species-compatible sequences.

Recent computational methods for protein binders use structure prediction^53^, diffusion models^31,33,54^, language models^32,55^, and physics-based modeling^56^ to propose sequences with high predicted affinity. However, high affinity alone is often insufficient in physiological contexts, particularly near membranes, where receptor topology, membrane curvature, and the glycocalyx create steric barriers that hinder productive binding. APPRAISE addresses this limitation by incorporating competitive AlphaFold-based complex modeling with rapid, physics-informed analyses of surface complementarity and steric accessibility. As a result, APPRAISE can rank peptide–receptor interactions in the context of capsid display and is especially suitable for designing ligands that remain geometrically accessible when presented on the bulky, icosahedral surface of AAV capsids for BBB targeting.

Because APPRAISE relies on competitive pairwise modeling within a peptide pool, it performs best when the pool includes at least one known or strong binder that provides a reference for ranking. In the absence of such positive binders, the model cannot reliably identify novel hits, limiting its utility for fully *de novo* discovery. EvoPRAISE overcomes these limitations by coupling APPRAISE with an iterative *in silico* directed evolution strategy. In each round, sequence diversity is introduced around current leads, for example through single-site saturation mutagenesis at selected positions, followed by competitive modeling with APPRAISE to select improved variants. By repeating this process, EvoPRAISE can discover high-affinity binders even without prior peptide knowledge, thereby expanding the accessible sequence space and reducing reliance on large animal cohorts that may not translate across species.

In this study, we demonstrate the utility of EvoPRAISE in the Syrian hamster. Capsids displaying EvoPRAISE-designed peptides were able to cross the BBB following intravenous administration and achieved widespread parenchymal transduction in the CNS. Within the CNS, we observed regionally enriched tropism in the cortex, hippocampus, thalamus, striatum, and rostral dorsal spinal cord. These regional differences are likely attributable to spatial heterogeneity in endothelial receptor expression, such as Ly6e, rather than to extensive post-entry diffusion. It is reported that post-entry diffusion is restricted by electrostatic interactions with extracellular matrix components, including perineuronal nets and heparan sulfate proteoglycans^57^. Along the evolutionary trajectory toward Cap-PF1.7, the peptide also underwent biochemical adaptation. Initial increases in hydrophobicity promoted interface formation and stabilization, while subsequent reintroduction of polarity restored solubility and conformational flexibility. A recurring pattern emerged, characterized by a hydrophobic core surrounded by a polar rim, which is typical of protein–protein interfaces. Oscillations in GRAVY values are consistent with exploratory adaptation and suggest stepwise testing of local compatibility with the receptor and surrounding structures before convergence on a functional optimum^58,59^. Such exploratory peptide evolution likely arose from the combination of saturation mutagenesis and progressive evolutionary algorithms, which promote sequence diversification and help prevent premature convergence on local optima in receptor binding.

Despite the remarkable success of EvoPRAISE there is still scope for improvement. Firstly, the performance of peptides obtained from a random input library can depend on library size. Very small libraries increase the risk of trapping at local optima on the binding score landscape. There is no field wide rule for a minimum starting size because the optimal choice depends on the fitness landscape and screening capacity. Researchers should also note that the sequence space explored by this iterative scheme grows quadratically with peptide length (*O(L*^2^), where *L* is the peptide length in amino acids), whereas the theoretical diversity of all possible peptide sequences increases exponentially (*O(*20*^L^)*). These results imply that the peptides obtained are unlikely to represent globally optimal solutions. In practical applications, several iterations of EvoPRAISE combined with experimental validation are needed to identify functionally robust candidates. The second issue with EvoPRAISE is that runtime is dominated by AlphaFold Multimer inference. Pairwise competition scales quadratically with the number of variants, which is impractical for very large libraries. For single site saturation per round, the modeled variants per round scale approximately linearly with peptide length, about 19 times *L* for one site scans. To address the computational cost of EvoPRAISE and to enable integration with experimental workflows, several complementary strategies can be applied. More efficient evolutionary search algorithms can be used to reduce computational expense, such as simple genetic algorithms with small populations, low mutation rates, and early stopping criteria to prevent unnecessary modeling. In addition, a compact candidate peptide binder library generated by EvoPRAISE can be screened *in vivo* using a focused design strategy. This approach is more practical for non-model species and can provide quantitative feedback for subsequent computational iterations. Finally, EvoPRAISE inherits uncertainties from predictive models and may not account for downstream barriers such as endocytosis, endosomal escape, intracellular trafficking, and uncoating. The development of reliable prediction methods and databases for identifying proteins suitable as AAV targets is highly desirable. Resources such as the Cell Surface Protein Atlas^60^, and brain vascular proteomic atlases^61^ provide a foundation for identifying potential AAV receptors. These datasets allow candidate proteins to be filtered by surface localization and endothelial enrichment, improving the selection of receptors that are accessible at the BBB for experimental validation.

In summary, EvoPRAISE couples competitive structure modeling with iterative *in silico*-directed evolution to convert initial peptide candidates into binders for specified targets. This methodology enables the design of AAV capsids without extensive prior knowledge and facilitates the identification of peptide ligands predicted to traverse the BBB and achieve CNS delivery across diverse species. Thus, experimental access to species-specific physiology in non-model animals is expanded. As such, EvoPRAISE advances rational capsid design toward clinically relevant BBB penetrant vectors.

## ACKNOWLEDGMENTS

This work was supported by the Japan Society for the Promotion of Science (JSPS) Grant-in-Aid for Early-Career Scientists (23K14130) (to H.O.), the JSPS Grant-in-Aid for Grant-in-Aid for Transformative Research Areas (A) (23H04941) (to G.A.S.), and the Suntory Rising Stars Encouragement Program in Life Sciences (to G.A.S.).

## AUTHOR CONTRIBUTIONS

H.O. designed the study. H.O. performed the animal experiments. H.O. and S.F. performed the histology analysis. H.O. and S.F. generated AAVs. H.O. performed the computational peptide design. H.O., wrote the original manuscript, which was reviewed and edited by G.A.S. G.A.S. supervised the study.

## INCLUSION AND DIVERSITY

We support inclusive, diverse, and equitable conduct of research.

## STAR★METHODS

### KEY RESOURCE TABLE

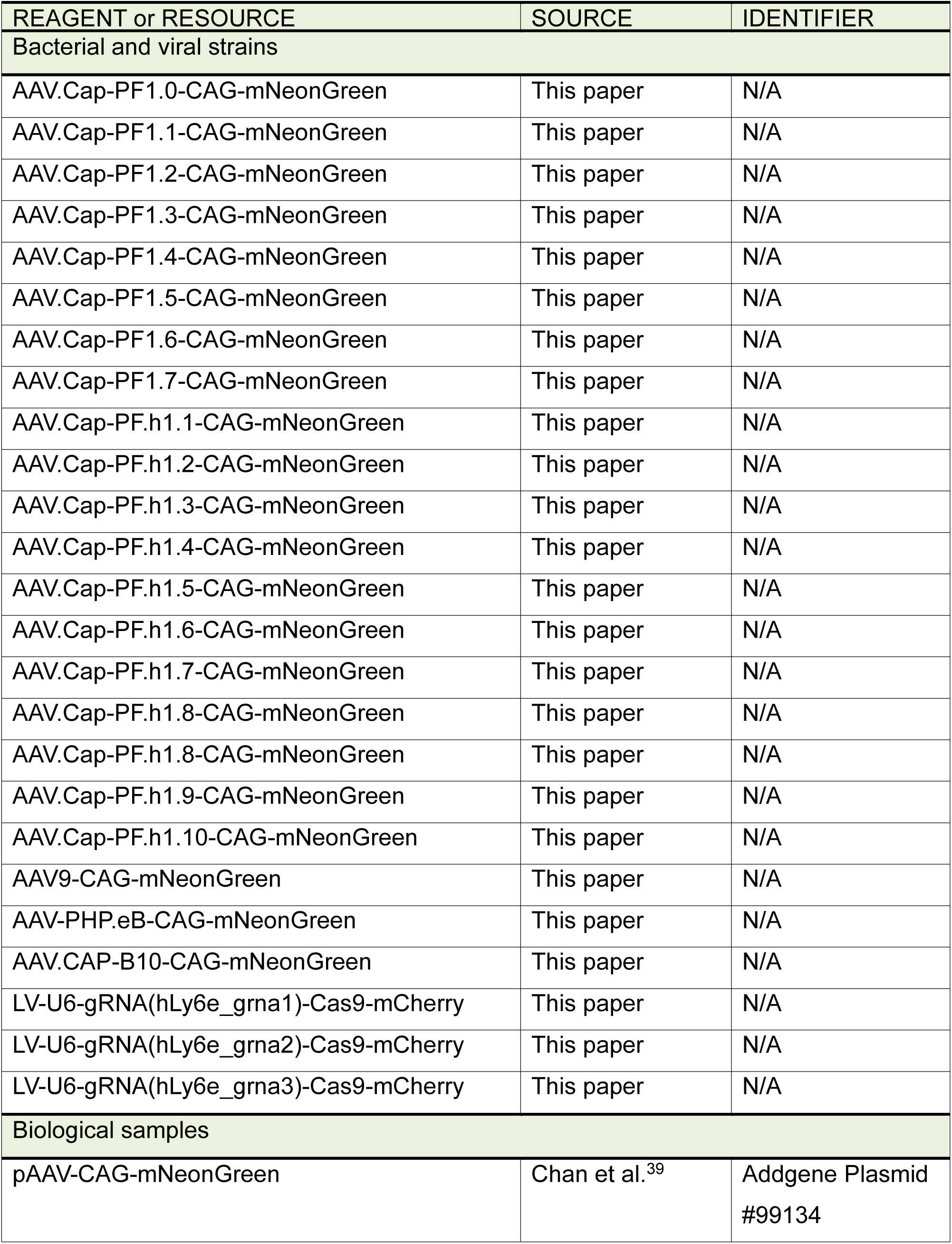
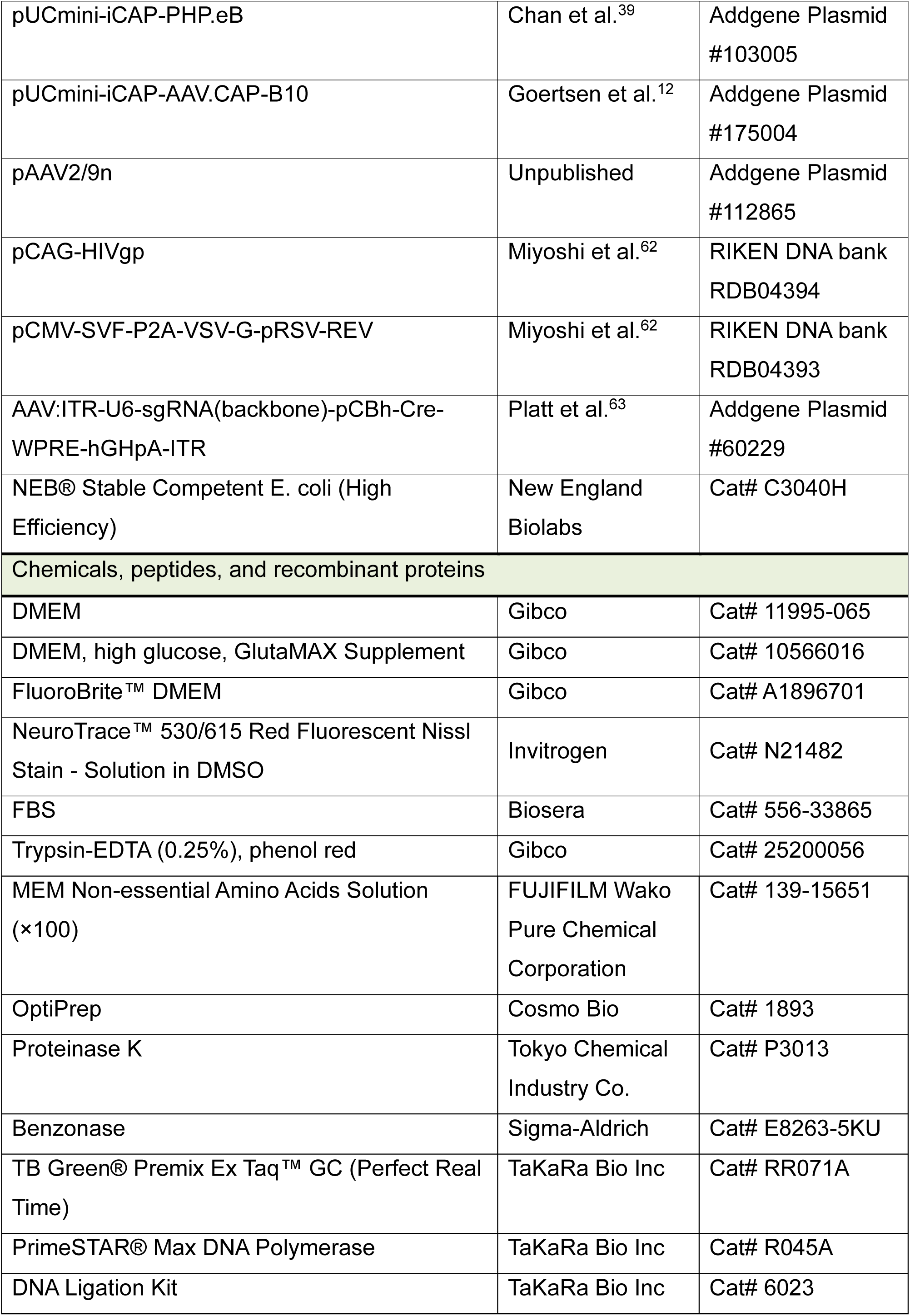
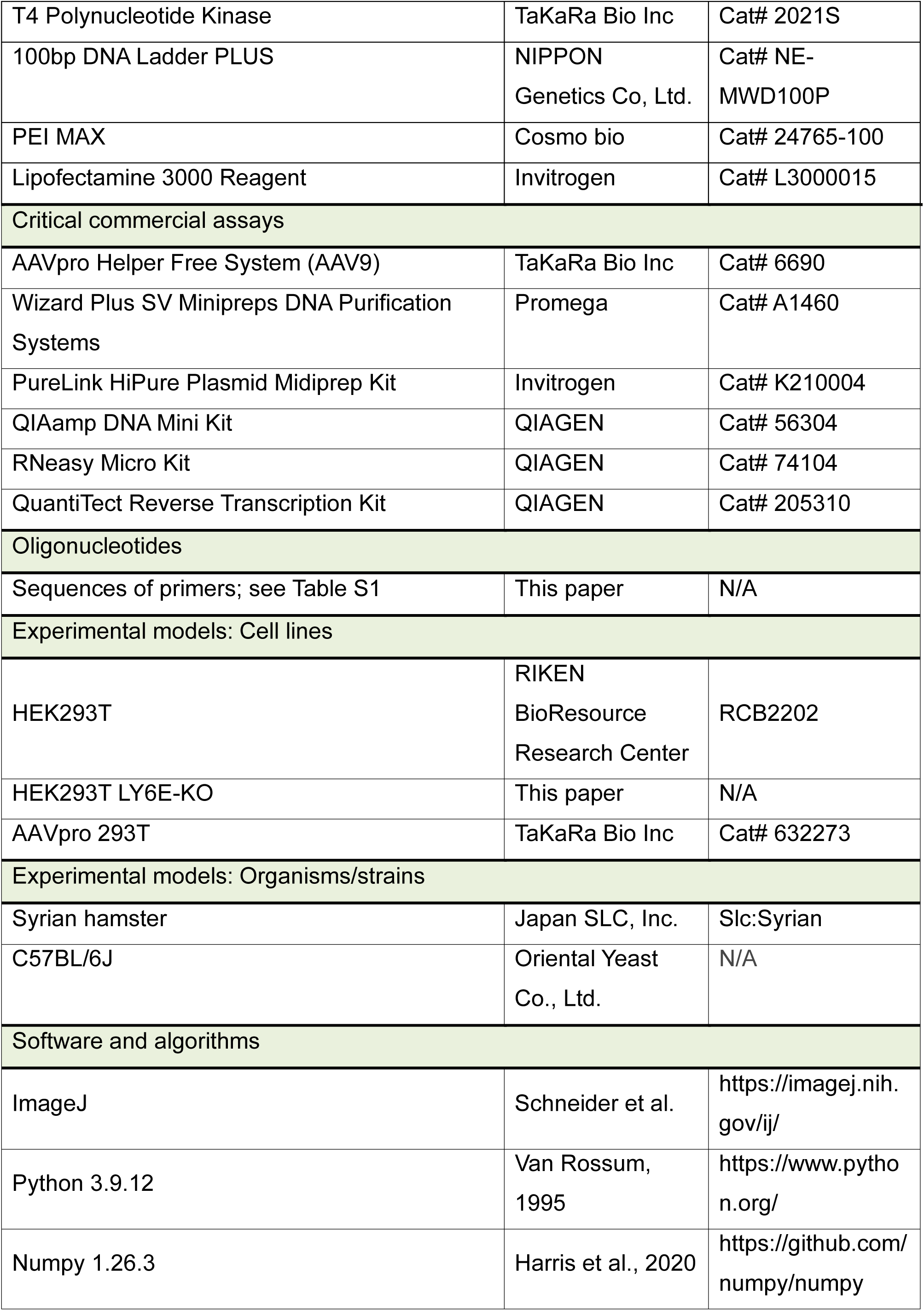
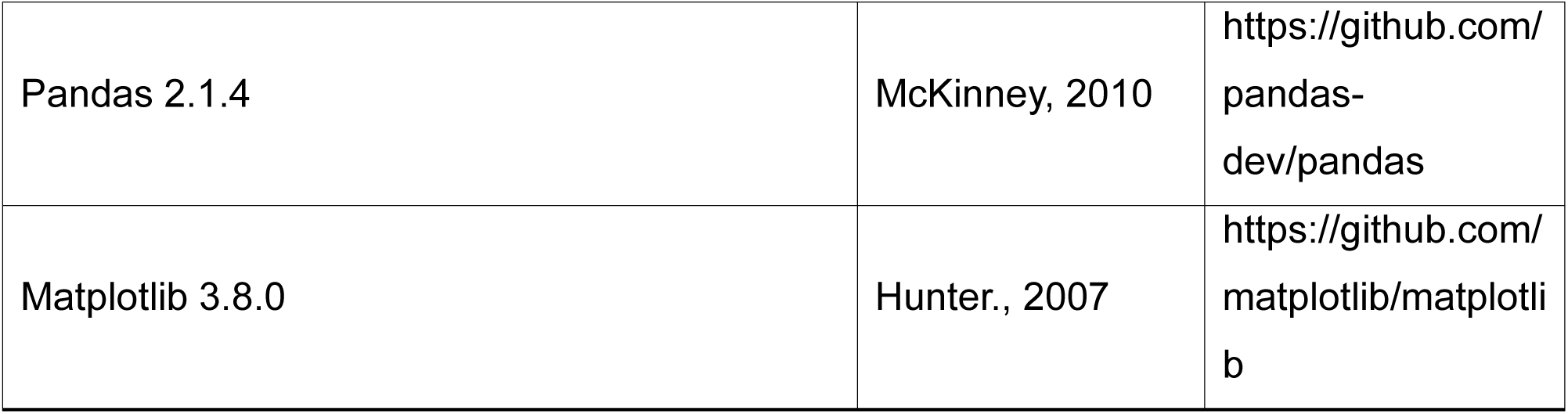

### RESOURCE AVAILABILITY

#### Lead contact

Further information and requests for resources and reagents should be directed to and fulfilled by the lead contact, Hiroaki Ono.

#### Material availability

All plasmids will be made available from the lead contact upon reasonable request.

#### Data and code availability

Any additional information required to reanalyze the data reported in this paper is available from the lead contact upon request.

### DECLARATION OF GENERATIVE AI AND AI-ASSISTED TECHNOLOGIES IN THE WRITING PROCESS

During the preparation of this work, the authors used ChatGPT in order to improve grammar and writing clarity. After using this tool/service, the authors reviewed and edited the content as needed and take full responsibility for the content of the publication.

### EXPERIMENTAL MODEL AND SUBJECT DETAILS

#### Evolutionary optimization combined with APPRAISE (EvoPRAISE) for AAVs

EvoPRAISE in this study comprised two processes: the identification of seed peptides and the evolution of these seed peptides. Custom Python code was used for the seed peptide identification process. In all, 100 variant peptides were generated by inserting a randomized sequence (7-mer) between AA588-589 (VP1 indices) of the AAV9 capsid. The APPRAISE method was employed to identify peptides that would serve as the seeds for directed evolution. Specifically, peptides with stronger affinity for the target sequence than AA587-594 of the AAV9 capsid (AQAQAQTG) and AA587-601 of PHP.eB (DGTLAVPFKAQAQTG) were identified. Affinity was evaluated using Delta_B as the metric, which is the default setting of the APPRAISE method. The chosen target sequences were the extracellular domains of hamster Ly6e (A0A1U7Q6V5_MESAU) (AA27-118) and human Ly6e (Q16553 · Ly6e_HUMAN) (AA21-112). The extracellular domains were determined as homologous sequences based on alignment with the AA27-110 peptide of mouse Ly6A (P05533 · LY6A_MOUSE), which was used in the demonstration of the APPRAISE method.

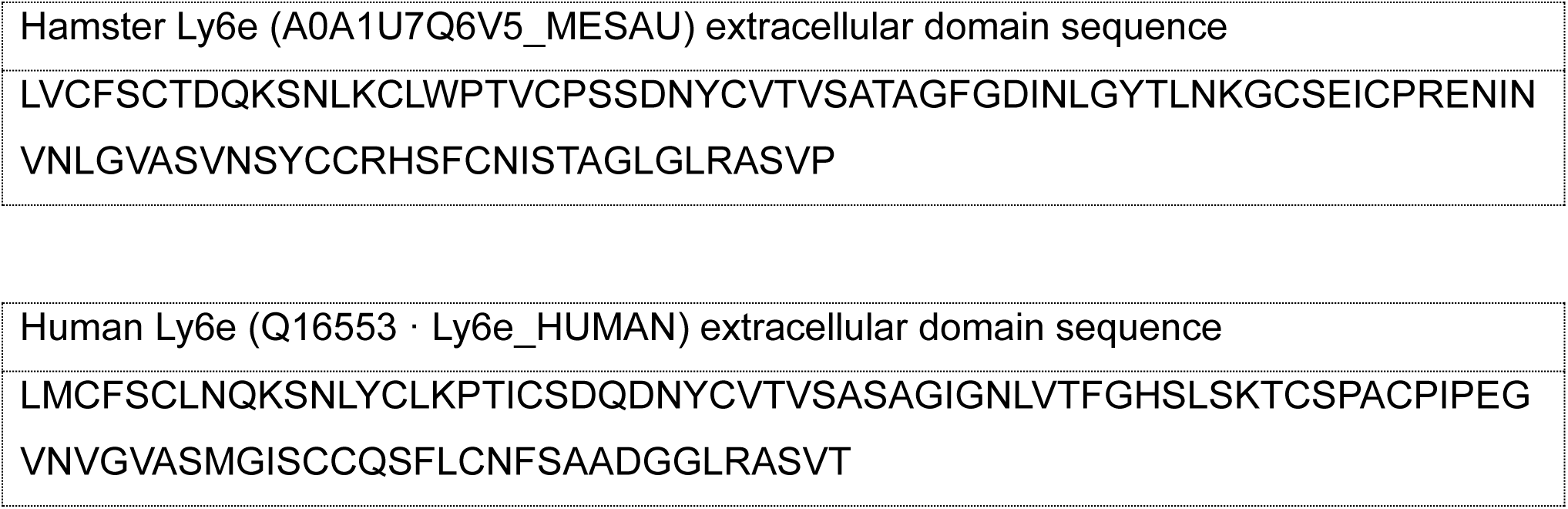

The publicly available version of APPRAISE was used (available at https://github.com/GradinaruLab/APPRAISE). Briefly, FASTA-format files containing the

amino acid sequence of the target receptor and peptide sequences were prepared for structural prediction. Batch-mode ColabFold (version 2.1.14) was used for complex modeling on Google Colaboratory equipped with an NVIDIA Tesla V100 SXM2 16 GB GPU or NVIDIA A100 Tensor Core 40 GB GPU. Unless otherwise specified, the AlphaFold model version alphafold-multimer-v2 was used as the default setting. Each model was recycled 3 times, and 5 models were generated per competition. Quantitative analyses of the resulting structures were performed in PyMOL (version 2.3.3) using custom scripts originally developed by Ding et al^37^. The following parameters were calculated: the number of peptide atoms at the receptor interface (*N* ^POI^ ; defined by an interatomic distance cutoff of 5 Å); the number of peptide atoms clashing with the receptor (*N* ^POI^ ; defined by a distance cutoff of 1 Å); the peptide binding angle (*θ*; defined as the angle between the vector from the receptor centroid to the anchor and the vector from the receptor centroid to the peptide centroid); and the peptide binding depth (*d*; defined as the difference between the distance from the receptor center to the closest peptide atom and the minor radius of the receptor ellipsoid hull, normalized by the minor radius). The minor radius of the receptor ellipsoid hull was calculated using HullRad 8.1, which yielded a value of 15.1 Å for Ly6e. Finally, the binding propensity metric *ΔB*^POI,competitor^ was computed as the difference in total binding scores between competing models. In the first round, the ranking was generated by comparing each peptide against the PHP.eB peptide as the reference competitor.

To perform peptide evolution, a site-saturation mutagenesis library was generated using custom scripts. Each amino acid residue in the selected peptide was substituted with all possible amino acids at that position. The APPRAISE pipeline, using the same parameter settings as described above, was then applied to evaluate and rank the peptide variants based on their predicted affinity for the target proteins. For *Round N* (0 ≤ *N* ≤ *L*, where *L* is the peptide length in amino acids), each mutant library was compared against the top-ranked peptide from *Round N - 1* to identify improved variants. In *Round N + 1*, the mutated position identified in *Round N* was fixed, and site-saturation mutagenesis performed on the remaining *L−N* residues. Thus, the evolutionary process for the 7 amino acid peptide entailed 7 iterative rounds of mutation. The final peptide obtained represented the product of *in silico* directed evolution. All complex structures in this study were modeled using template based modeling settings.

msa_mode = "MMseqs2 (UniRef+Environmental)" num_models = 5

num_recycles = 20

stop_at_score = 100 use_custom_msa = False use_amber = False use_templates = True

model_type = "auto" or "alphafold2_multimer_v2"

#### Plasmid construction

All capsid variants were generated using a common mutagenesis strategy. First, the entire AAV9 capsid coding sequence was subcloned into the multiple cloning site of the pUC19 vector (pUC19-Cap9). The amino acid sequences of the peptide binders were converted into nucleotide sequences optimized for human codon usage. The optimized peptide-coding sequences were inserted into the peptide insertion site of pUC19-Cap9, located between residues 588 and 589 of the AAV9 capsid (VP1 indices), by vector PCR using PrimeStar Max DNA polymerase followed by blunt-end ligation. The mutated capsid fragments were amplified using primers flanking the mutagenized region and cloned into the corresponding site of the pUCmini-iCAP-PHP.eB (Addgene plasmid #103005). Sequences of all the oligonucleotide primers used for plasmid construction are given in Table S1.

#### AAV preparation

The protocol for AAV production shown below was based on that described by Challis, et al., 2019^64^ with modifications. AAVpro 293T was cultured in 150 mm dishes in a culture medium containing DMEM (high glucose) (Gibco), 10% (v/v) fetal bovine serum (FBS), and 1% penicillin-streptomycin (PS) at 37°C and 5% CO2. pAAV, capsid of interest, and pHelper plasmid were transfected into cells, reaching 90-100% confluency. A pAAV: capsid: pHelper plasmid ratio of 1:4:2 based on micrograms of DNA (i.e., 5.7 μg of pAAV, 22.8 μg of capsid, and 11.4 μg of pHelper) was used for the transfection. On the day following transfection, the culture medium was exchanged for a 20 mL culture medium containing DMEM (high glucose, GlutaMAX) (Gibco), 5% (v/v) FBS, 1% MEM Non-Essential Amino Acids solution (NEAA) (Wako), and 1% PS. After 72 h following transfection, the culture medium was collected and exchanged for 20 mL of fresh culture medium containing DMEM (high glucose, GlutaMAX), 5% (v/v) FBS, 1% MEM NEAA, and 1% PS. The collected culture medium was stored at 4°C. Cells were subsequently collected using a cell scraper 144 h after transfection. The collected culture medium obtained 72 h post-transfection was separated into supernatant and cell pellet by centrifugation (2000 x g, 20 min). Polyethylene glycol was added to the supernatant to a final concentration of 8%. The resulting solution was then incubated on ice for 2 h. The solution was centrifuged (5000 x g, 30 min) and the pellet suspended into 2 mL Tris MgCl2 (10 mM Tris pH 8.0, 2 mM MgCl2). The suspension was subjected to three rounds of freeze-thaw cycles. Benzonase (100 U/mL) was then added and the mixture incubated at 37°C for 1 h. The supernatant solution was mixed with the cell solution, and the resulting mixture centrifuged (2,000 x g, 10 min) to obtain the supernatant. Iodixanol density gradient solutions (15%, 25%, 40%, and 60% (wt/vol)) were prepared and poured into an Optiseal centrifuge tube (Beckman). Tubes were transferred to a Type 70 Ti rotor (Beckman) and centrifuged at 350,000 x g for 2 h 25 min. In each case the 40% iodixanol layer containing the virus was collected. The purified viral solution was obtained by ultrafiltration using an Amicon filter device (Merck) and resuspension into Dulbecco’s phosphate-buffered saline.

For AAV titration, purified virus solution was treated with Benzonase (0.05 U/mL, 37°C, 1 h) followed by Proteinase K (0.25 mg/ul, 37°C, 1 h). The viral genome was subsequently obtained by phenol-chloroform-isoamyl alcohol extraction followed by isopropanol precipitation. The titer (v.g./mL) of the AAV was calculated by quantifying the number of WPRE sequences in the sample using qPCR. pAAV-CAG-mNeonGreen (Addgene Plasmid #99134) was used as the standard. The qPCR protocol was as follows: 95°C for 60 s (initial denaturation) followed by 45 cycles of 95°C for 10 s, 60°C for 30 s using TB Green® Premix Ex Taq™ GC (TaKaRa Bio Inc). The final purified virus samples were stored at -80°C. Titers of recombinant AAV vectors were determined by quantitative PCR.

#### AAV vector administration, tissue processing and imaging

In the hamster experiments, AAV vectors were administered intravenously via retro-orbital injection at doses of 1-5 × 10^13^ v.g. as indicated in the figures and corresponding legends. In the mouse experiments, AAV vectors were administered intravenously via retro- orbital injection at doses of 1 × 10^12^ v.g. After 4 weeks of expression, hamsters and mice were anesthetized with isoflurane and transcardially perfused with approximately 50 mL of

10% sucrose and then another 50 mL of 4% paraformaldehyde (PFA) in 0.1 M phosphate buffer saline pH 7.4 (PBS). Organs were post-fixed overnight in 4% PFA at 4°C, incubated overnight in 30% sucrose in 0.1 M PBS at 4°C. Finally, organs were cut into 50 μm sections using a Leica CM1950 Cryostat. Images were acquired with a fluorescence microscope (Axio Observer 7, ZEISS) and processed using ZEN 3.4 (Zeiss) and ImageJ software.

#### Lentivirus production

Lentivirus production was performed according to RIKEN’s protocol (https://dnaconda.riken.jp/Form_PDF/lntPrepen.pdf) with some modifications. HEK293T cells were seeded onto six 15 cm culture dishes (Falcon) at 1.69 × 10^7^ cells/dish. After 24 h incubation, the medium was changed to 19 mL Improved Minimum Essential Medium (IMEM) supplemented with 15 µM chloroquine (Wako) without PS and FBS. Cells were then incubated for an extra 1 h at 37°C, 5% CO2. Transfection was conducted using the calcium phosphate precipitation method. In brief, the plasmid mixture was prepared by mixing 281 µg target plasmid, 165 µg pCAG-HIVgp, and 165 µg pCMV-SVF-P2A-VSV-G-pRSV-REV, and made up to 7.5 mL with double distilled water. Next, 500 µL of 2.5 M CaCl2 (Wako) was applied to the bottom of the tube containing this plasmid mixture and vortexed vigorously for 10 s. An 8 mL aliquot of borate buffered saline (Wako) buffer was added dropwise to the plasmid mixture while gently mixing. After incubation at room temperature for 8 min, 2.7 mL of the plasmid mixture was applied to the HEK293T cells and the mixture incubated at 37°C, 5% CO2 for 1 h. 10 % FBS was then added to medium. After incubating for 13 h, medium was changed to 15 mL IMEM containing 1.5 mM sodium butyrate (Wako) and 1.5 mM caffeine (Tokyo Chemical Industry). Following further incubation at 37°C, 5% CO2 for 34 h, the supernatants were collected and centrifuged (4°C, 900 g, 10 min) to remove cell debris. Finally, the supernatant was filtered through a 0.45 μm PES filter unit (Merck Millipore), divided into three 30 mL tubes (Beckman Coulter, 326823) by layering onto 10 mL of 10% sucrose solution, and centrifuged at 4°C, 15,000 rpm for 4 h in an ultracentrifuge (Beckman Coulter) to pellet the virus. The resulting pellet was vigorously resuspended in 3 mL of chilled PBS and 10 µL aliquots of the suspension were stored at -80°C.

Titer evaluation was performed to determine the infectious titer. HEK293T cells were seeded into 24-well plates (Corning) at 3.53 × 10^4^ cells/well. After 4 h of incubation, serially diluted virus solutions (0, 1,000, 2,000, 4,000, 8,000, 16,000, 32,000-fold dilution) were applied to the cells in a final volume of 0.5 mL per well, followed by incubation at 37°C in 5% CO₂ for an additional 72 h. Cells were then detached with 0.25% trypsin-EDTA (Wako) and analyzed using Cell Sorter MA900 to quantify the percentage of fluorescent protein-positive cells. The infectious titer unit (IFU/mL) was calculated using the following formula:

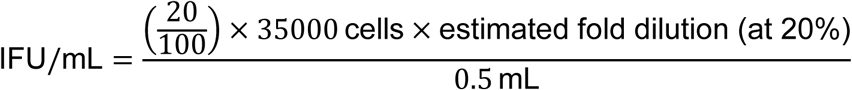

where 0.5 mL corresponds to the volume of virus solution applied per well. The estimated fold dilution at 20% was determined by approximate linearization (Microsoft Excel) based on infection efficiency values below 20%.

#### Establishment of the LY6E-KO cell line

CRISPR-Cas9–mediated knockout of *LY6E* was performed using a lentiviral delivery system. A guide RNA (gRNA) was selected using CRISPRdirect (https://crispr.dbcls.jp/) with the following sequence.

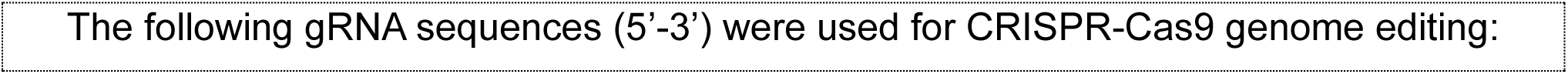
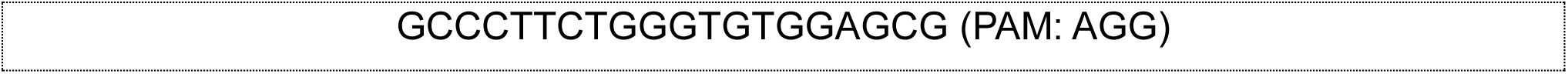

The gRNA was inserted into the U6-sgRNA backbone derived from AAV:ITR-U6- sgRNA(backbone)-pCBh-Cre-WPRE-hGHpA-ITR (Addgene plasmid #60229). For lentiviral backbone construction, Lenti-Cas9-blast (Addgene plasmid #52962) was modified by inserting mCherry between the BamHI and EcoRV restriction recognition sites, generating Lenti-Cas9-mCherry. Subsequently, the U6-sgRNA cassette was introduced at the upstream NheI site of the Cas9-mCherry expression cassette, yielding Lenti-U6-gRNA-Cas9-mCherry. Lentiviral particles were produced from this vector and applied to HEK293T cells at a multiplicity of infection corresponding to 1. One week after infection, mCherry-positive cells were single-cell sorted using a Cell Sorter MA900 (Sony). Clonal populations were expanded, and genomic DNA was extracted using QIAamp DNA Mini Kit (QIAGEN). The genomic region flanking the gRNA target site was amplified by PCR and subjected to Sanger sequencing for genotyping, confirming frameshift mutations in *LY6E*.

#### Cell culture for AAV infectivity evaluation

HEK293T cells were seeded at 80% confluency in 96-well plates and maintained in DMEM supplemented with 5% FBS, and PS (100 U/mL) at 37°C in 5% CO2. Target candidates (150 ng) were transiently expressed in HEK293T cells. DNA transfection was performed using Lipofectamine 3000 according to the manufacturer’s instructions (Invitrogen). Cells expressing each target candidate were transduced with engineered AAV variants at 5 × 10^9^ v.g. per well in triplicate 24 h after transfection. Receptor-expressing cells were transferred to 96-well plates at 20% confluency and maintained in FluoroBrite DMEM supplemented with 0.5% FBS, 1% NEAA, PS (100 U/mL) at 37°C in 5% CO2. Images were acquired with a fluorescence microscope (Axio Observer 7, ZEISS) and processed using ZEN 3.4 (Zeiss) and ImageJ software.

#### Cell culture fluorescence image quantitation

All image analyses were performed using a custom Python-based image processing pipeline developed for automated quantification of transduction efficiency and fluorescence intensity. The pipeline processed paired bright-field and fluorescence images, extracting metrics including total cell area, transduced area, transduction percentage, and fluorescence intensity per transduced area. For both image types, background subtraction was performed by first converting RGB images to grayscale, followed by Gaussian blurring (skimage.filters.gaussian) and subtracting the blurred image from the original. For bright-field images, a Gaussian filter with sigma=30 and truncate=0.35 was applied. Histogram-based thresholding (skimage.filters.threshold_otsu) was then used on both the subtracted image and its inverted version to detect bright and dark regions corresponding to cell edges. These binary masks were combined and morphologically closed (skimage.morphology.closing with a disk of radius 2) to fill in cellular regions. The total cell area was determined by summing the pixels in this final mask. For fluorescence signal images, a Gaussian filter with sigma = 100 and truncate = 0.35 was applied, and the resulting blurred image was subtracted from the original to remove background fluctuations. Histogram-based thresholding was used to identify regions with strong fluorescence, corresponding to transduced cells. Small noise artifacts were removed using skimage.morphology.remove_small_objects (minimum size = 5). The transduced area was calculated as the total number of pixels in this cleaned mask. To quantify signal intensity, the fluorescence image was multiplied by the binary mask of transduced cells, and the resulting pixel intensities were summed to obtain the total brightness of transduced regions. This value was then normalized by transduced area to yield the brightness per transduced area. To avoid overestimation in cases with extremely low signal or cell area, a lower threshold of 0.1% was enforced for the transduction percentage. Paired bright-field and fluorescence images were automatically matched and analyzed from a specified folder. The image filenames were parsed to extract metadata such as serotype and dose. All calculated values were stored in a summary CSV file using the pandas library.

#### Fluorescence staining and image-based quantification of double-positive cells

To analyze the distribution of cells co-expressing two molecular markers, brain sections from the thalamus, cortex, and hippocampus were subjected to fluorescent staining and quantitative image analysis. Coronal brain sections of 30 μm thickness were mounted onto glass slides and air-dried overnight at room temperature. Sections were then briefly washed in PBS for 5 min, followed by a 10 min wash in PBST (PBS containing 0.1% Triton X-100), and two additional 5 min washes in PBS. Staining was performed by applying 200 μL of the 1/20 diluted NeuroTrace™ 530/615 Red Fluorescent Nissl Stain (Thermo Fisher Scientific) solution to each section and incubating at room temperature for 20 min. After staining, sections were washed again in PBST for 10 min and twice more in PBS (2 x 5 min). A final wash in PBS was performed either for 2 h at room temperature or overnight at 4°C. After washing, sections were coverslipped and stored at 4°C until imaging. Fluorescent images of each region were acquired using a fluorescence microscope. For each sample, two grayscale images representing different fluorescent markers were exported as 16-bit TIFF files. These images were analyzed using a custom Python script based on OpenCV, scikit-image, and matplotlib libraries. Each image was converted to grayscale if it contained RGB information, then binarized using fixed intensity thresholds selected empirically to minimize background noise while retaining cellular signals. In the representative analysis, a threshold of 100 was applied to the first image (channel 1) and a threshold of 10 to the second image (channel 2). Binarized images were processed using an 8-connected component labeling algorithm to identify signal-positive regions. These labeled regions, corresponding to individual cells, were characterized using morphological feature extraction. To assess signal co-localization, each cell identified in the first image was examined for pixel-wise overlap with the labeled regions in the second image. A cell was classified as double-positive if at least one of its pixels overlapped with a region in the second channel. The proportion of double-positive cells was calculated as the number of overlapping cells divided by the total number of cells in the channel that contained fewer cells. This procedure prevented overestimation due to differences in segmentation sensitivity between channels. The results were visualized as pie charts, with double-positive cells categorized as “Neuron” and the remaining single-positive cells as “Glia.” All analyses and visualizations were saved as image files corresponding to the specific brain regions.

#### Structural analyses

Peptide–receptor structures were modeled using AlphaFold-Multimer, which is based on AlphaFold3^29,35^. As the query sequence, we used the extracellular domain of Hamster Ly6e (UniProt ID: A0A1U7Q6V5_MESAU), as described earlier. All other parameters were kept at their default settings. AAV trimer–receptor complex models were constructed using a hybrid structural modeling approach inspired by the strategy described in Shay et al., 2023^38^. Trimers located at the AAV threefold symmetry axis were selected as the minimal binding interface that could approximate interactions between the full capsid and the putative receptor while maintaining computational tractability.

First, a peptide–receptor complex was modeled by inputting a 15-amino-acid peptide (residues 587–594 based on WT AAV9 VP1 numbering) from the AAV.Cap-PF1.7 variant together with the target receptor, as described above. Separately, a trimer model of the AAV.Cap-PF1.7 was generated using AlphaFold-Multimer. From the peptide–receptor model, the two residues with the highest predicted confidence (pLDDT score) Trp1’ and Met2’ were structurally aligned to the corresponding residues on the first subunit of the trimer model (coarse combined model). The two loops between Ala587 and Ala589 (VP1 indices) were remodeled using RosettaRemodel^44^ from the Rosetta software bundle (release 2018.48.60516). Lastly, these remodeled loops were merged to generate the final model. pLDDT scores for individual residues from the original AlphaFold-Multimer outputs were used to color images of the final model to generate a visual representation of predicted confidence. The final model was further optimized using the Rosetta fast-relax protocol.

#### RT-qPCR analysis of hamster brain

Total RNA was isolated from hamster brain tissue using the RNeasy Mini Kit (QIAGEN) according to the manufacturer’s instructions. Approximately 20–30 mg of brain tissue was rapidly frozen in liquid nitrogen, pulverized to a fine powder, and subsequently homogenized with a Microtube Homogenizer G10 (Bio Medical Science Inc.). cDNA was synthesized from total RNA using a QuantiTect Reverse Transcription Kit (QIAGEN), following the manufacturer’s protocol. GAPDH was used as the internal control. All oligonucleotide primer sequences used for qPCR are given in Table S1. Quantitative PCR was performed with TB Green® Premix Ex Taq™ GC (Perfect Real Time, TaKaRa Bio Inc) under the following conditions: initial denaturation at 95°C for 60 s, followed by 45 cycles of denaturation at 95°C for 10 s and annealing/extension at 60°C for 30 s. Relative gene expression levels were calculated using the 2^-ΔCt^ method, normalized to GAPDH. Data are grouped by gene. Mean ± standard deviation (SD) values are reported.

### QUANTIFICATION AND STATISTICAL ANALYSES

Data representation and statistical analyses were performed in python. One-way ANOVA followed by Tukey’s post hoc test was used in the analysis of RT-qPCR data. Statistical significance is indicated as follows: p < 0.05 (*), p < 0.01 (**), p < 0.001 (***), and p < 0.0001 (****). For comparisons between Ly6e and other proteins within each brain region, the Mann–Whitney U test was used, followed by multiple testing correction with the Benjamini–Hochberg method. Statistical significance is indicated as follows: p < 0.05 (*), p < 0.01 (**), p < 0.001 (***).

## SUPPLEMENTAL INFORMATION

**Figure S1.**
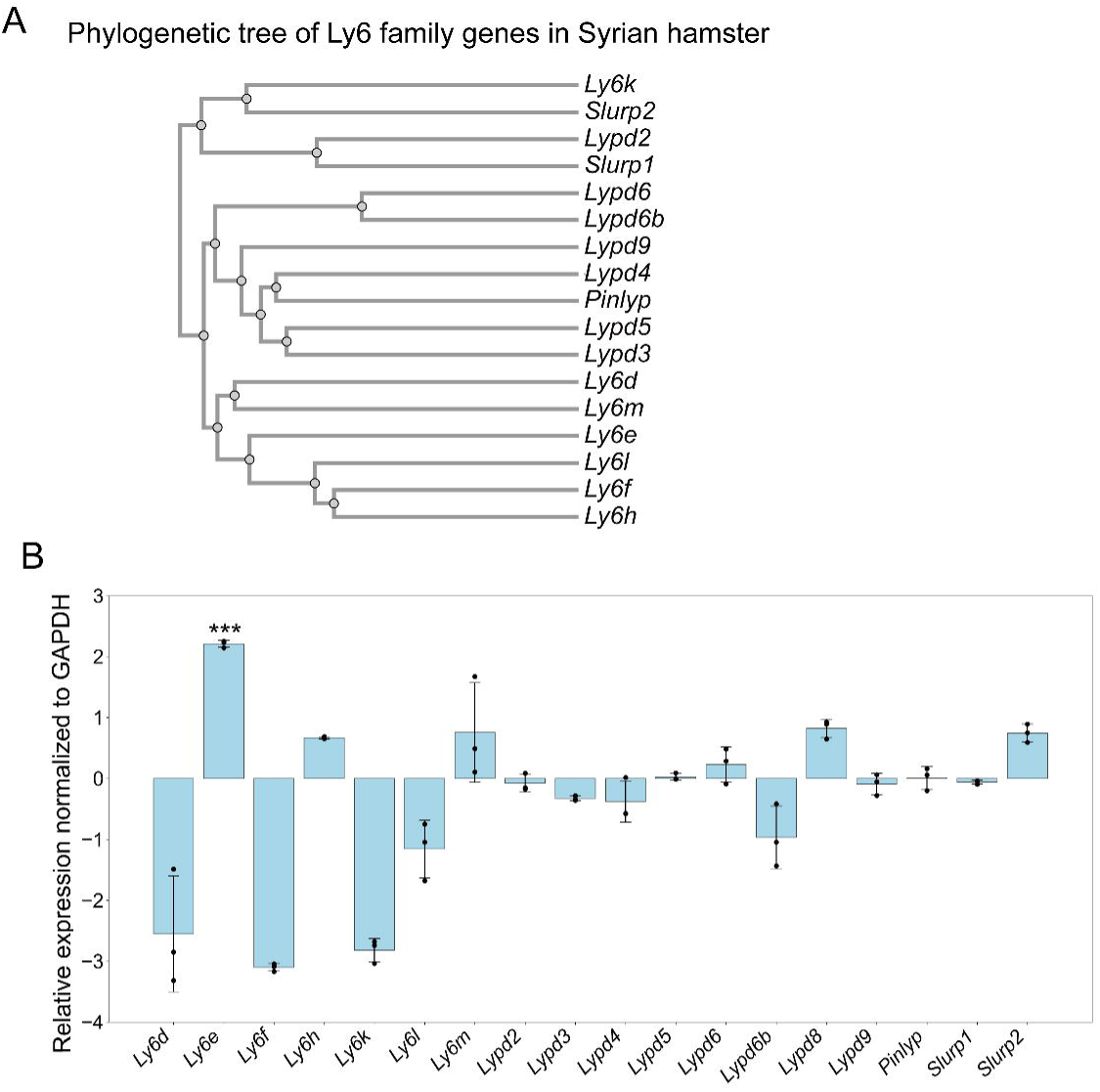
RT-qPCR analysis of Ly6 family gene expression in hamster brain (A) Ly6 family genes were manually extracted from the annotated genes in the golden hamster genome (Baylor, 2021) registered in the genome browser. A phylogenetic tree was constructed using Clustal Omega. (B) Expression levels of each Ly6 family gene were normalized. Data represent mean ± SD of n = 3 technical replicates. One-way ANOVA revealed significant differences among genes. Post hoc Tukey’s test indicated that Ly6e expression was significantly higher than that of most other Ly6 family members.

**Figure S2.**
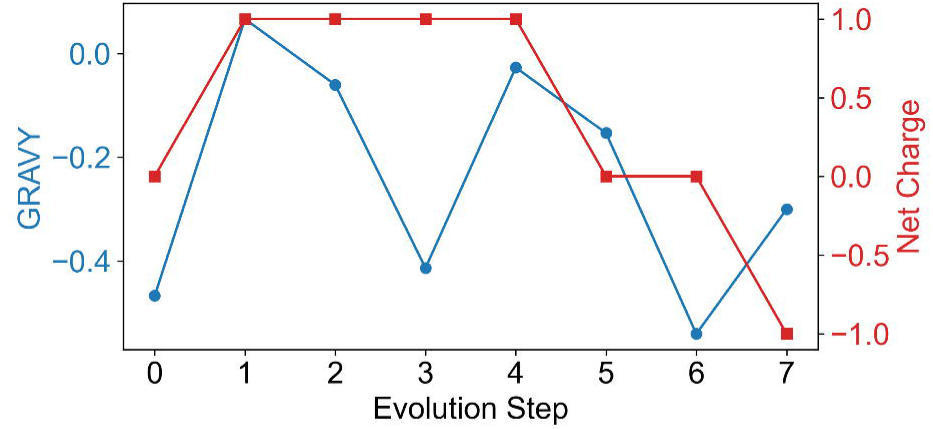
Cyclic changes in hydropathy and charge during sequence evolution. Average hydropathy (GRAVY score; blue line, left y-axis) and net charge (red line, right y-axis) were calculated for each sequence across the evolutionary steps. Hydropathy values were obtained by averaging Kyte–Doolittle indices across all residues in a sequence, whereas net charge was estimated at pH 7 by assigning Asp/Glu = –1, Lys/Arg = +1, and His = +0.1. A recurring pattern was observed in which an increase in sequence hydropathy was followed by compensatory adjustments in net charge during the evolutionary process.

**Figure S3.**
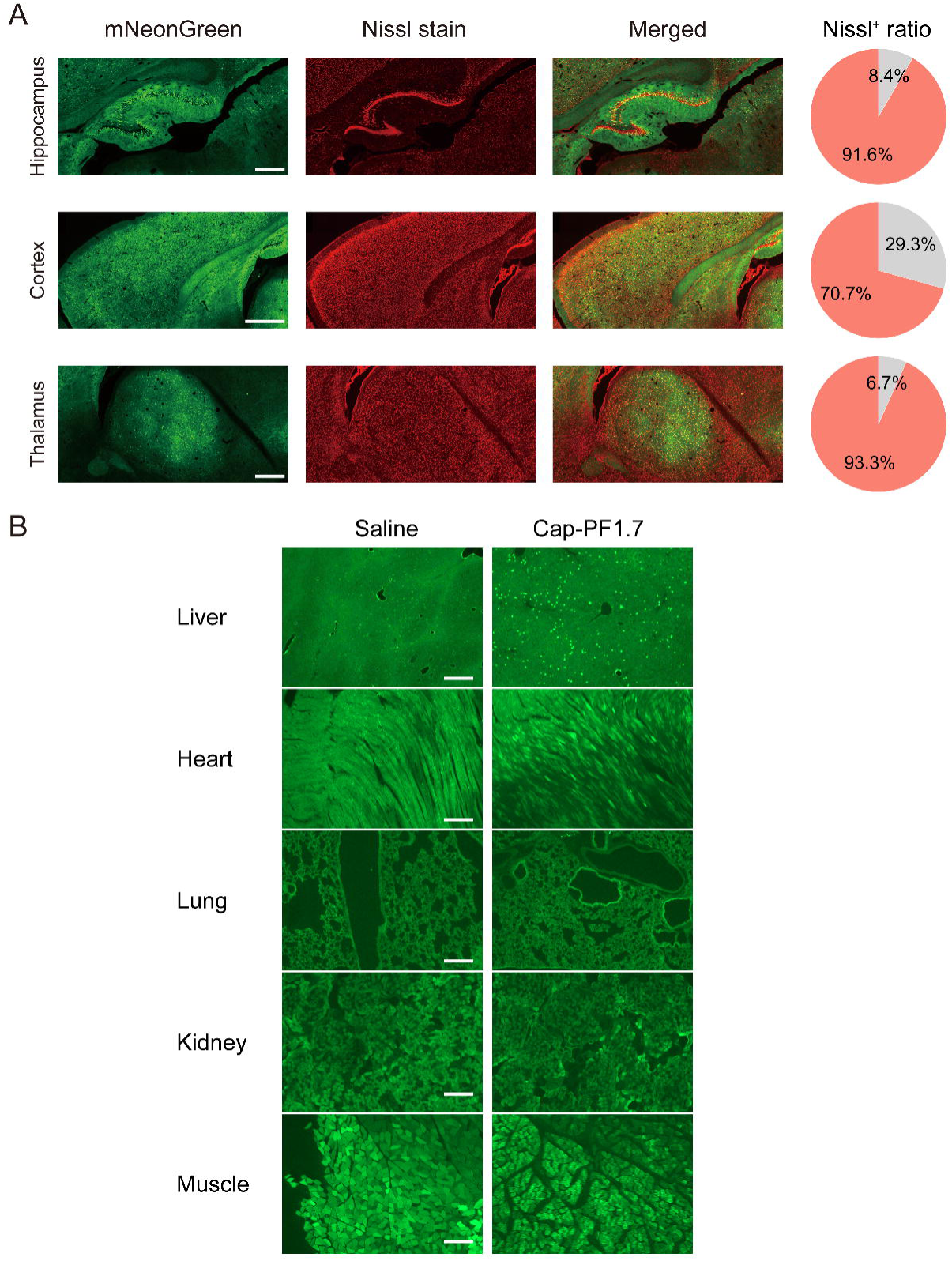
Cap-PF1.7 targeting cell type and off-target tissue validation (A) NeuroTrace assessment of representative brain regions to evaluate the cell types targeted by Cap-PF1.7. Colocalization of mNeonGreen fluorescence (driven by the CAG promoter) and NeuroTrace signal were assessed for each brain section. Pie charts indicate the proportion of NeuroTrace-positive cells among mNeonGreen-expressing cells. AAVs were administered intravenously to Syrian hamsters at a dose of 1 × 10^12^ v.g. per animal (n = 3 per condition). Four weeks after administration, transgene expression was assessed by visualizing mNeonGreen fluorescence throughout the brain. Scale bars, 500 µm. (B) Evaluation of mNeonGreen fluorescence (driven by the CAG promoter) in peripheral tissues. AAVs were administered intravenously to Syrian hamsters at a dose of 1 × 10^12^ v.g. per animal (n = 3 per condition). Four weeks after administration, transgene expression was assessed by mNeonGreen fluorescence. Scale bars, 500 µm.

Table S1. Sequences of oligonucleotides used for plasmid construction, RT-qPCR and cloning

